# Fluorescently-labeled split-QF hemidrivers: simplifying and enhancing methods enabling intersectional targeting of discrete cell types

**DOI:** 10.64898/2026.06.01.729389

**Authors:** Lydia G. Sanders, Grant Kroschell, Anneliese Ceisel, Caroline E. Clouatre, Yiqi Gao, Gianna Graziano, Diego Alfaro Carcoba, Claire Deng, Catalina Rodriguez, James H. Thierer, Jeff S. Mumm

## Abstract

Single cell transcriptomics data predict greater cell type diversity than previously appreciated, defined by combinatorial gene expression “codes”. These codes can enable intersectional targeting strategies for selectively labeling and manipulating predicted cell types, thereby facilitating functional tests to resolve cell-specific roles and refine cell-type definitions. One such approach cleaves driver components of binary expression systems (e.g., Gal4/UAS and QF/QUAS) into two co-dependent halves, an Activation Domain (AD) and a DNA Binding Domain (DBD). The transcriptional activity of these “split-drivers” (aka, hemidrivers) is only reconstituted in cells that co-express them. While widely used in invertebrates, these systems have yet to be systematically deployed in vertebrates. Here, we developed a series of fluorescently labeled split-QF and QF2 hemidrivers via direct reporter fusions or the 2A viral peptide co-expression system. Fluorescent labeling serves to 1) simplify hemidriver transgenic line creation and maintenance, 2) allow AD and DBD intersects to be visualized directly, and 3) enable robust quality control assessments of on-target efficacy and off-target activity. These resources can enhance transgenic targeting specificity, enabling functional dissections of cell types revealed by single cell transcriptomics.

**SUMMARY:** We engineered fluorescently labeled split-QF and -QF2 hemidrivers in zebrafish to simplify intersectional targeting tool generation and enable robust quality control assessments of split-driver function.

## INTRODUCTION

Our understanding of cellular diversity has been revolutionized by single cell transcriptomics, enabling the prediction of many transcriptionally distinct, and possibly functionally distinct, novel cell types and subtypes. Methods for exclusively targeting these newly defined cell types are needed to test functional roles to advance understanding of how complex tissues, such as the brain, are organized. Single-cell RNA sequencing (scRNAseq) datasets provide candidate marker genes for targeting putative cell subtypes^1–4^ through direct integration of transgenes at defined loci. However, while generally adequate for labeling major cell classes, a single marker gene is typically not sufficient for selective targeting of discrete transcriptomically-defined cell types and subtypes.

To enhance specificity, intersectional targeting systems have been developed that restrict reporter/effector transgene expression to the subset of cells that co-express two or more marker genes. For instance, Gal4 and QF transcriptional drivers, or “holodrivers”, can be split into two requisite “hemidriver” halves, the transcriptional activation domain (AD) and the DNA-binding domain (DBD)^5–8^. The addition of complementary dimerization domains to each allows a functional holodriver to be reconstituted only in the cells that co-express the AD and DBD hemidrivers. Activation of UAS- and QUAS-based “reporters/effectors” is thereby restricted to the intersection of the AD and DBD expression domains, allowing cell types and subtypes defined by the co-expression of two genes to be selectively labeled and/or manipulated for functional assays.

Gal4 and QF hemidrivers have been used to dissect neuronal subtype functions in *Drosophila*^9^ and *C. elegans*^10–12^. Additionally, an intersectional system based on co-expression of Cre recombinase and QF has been used to target and define functions for neuronal subtypes in the zebrafish retina^1^. However, these systems require specially designed responder lines containing loxP Cre binding sites and are therefore not immediately compatible with most existing UAS/QUAS lines. Although split Gal4 hemidrivers have been shown to be operable in zebrafish^13^, split driver systems have yet to be widely utilized in a vertebrate model system. While the Gal4/UAS system exhibits DNA methylation-based epigenetic silencing due to the CpG-rich UAS^14,15^, the QF/QUAS system has been optimized to decrease CpG dinucleotides and subsequently appears to be less susceptible to this issue^16^. We therefore chose to focus on the QF/QUAS system for an initial effort toward generating new intersectional targeting resources. Previously, usage of QF with broadly expressed reporters was found to have lethality issues in *Drosophila*^17^. This prompted the creation of QF2, a QF driver lacking the Middle Domain (MD), which was found to be a major source of toxicity^18^. In addition to reduced lethality and developmental effects, QF2 was shown to result in similar and/or slightly higher QUAS responder activation in *Drosophila*^18^. Accordingly, we chose to explore and compare the use of both QF and QF2 in generating these resources.

Successfully adapting a split driver system to zebrafish will facilitate two- and even three-gene intersectional targeting. Achieving multifactorial targeting is particularly important for studies of neuronal subtype function as highlighted in recent reports linking functional, transcriptional, and anatomical identities of neurons in the zebrafish retina^1,4,19^ and brain^3,20,21^. More recently, using WARP (for <u>W</u>hole-brain neuronal <u>A</u>ctivity and <u>R</u>NA <u>P</u>rofiling) to profile neurons responding to eight visual stimuli in the larval zebrafish brain, more than 2,000 functionally-genetically-anatomically neuronal subpopulations were predicted^22^. Importantly, of the 332 combinations used to genetically delineate these neuronal subtypes, 92% can be targeted using three or fewer marker genes, underscoring the value of adapting additional intersectional targeting methods to vertebrates.

Here, we present a fluorescent reporter-labeled split-QF hemidriver system developed in zebrafish for enhancing the expression specificity of QUAS-encoded reporter/effector transgenes. Reporter labeling simplifies transgenic line creation, facilitates direct visualization of AD and DBD expression overlap (i.e., targeting), and allows quantification of transactivation efficiency relative to AD and DBD expression levels and across hemidriver variants. We identified optimized topologies for two hemidriver labeling options, direct reporter fusions and 2A viral peptide-based co-expression. This study highlights the importance of correlating DBD and AD expression levels relative to each other and to transcriptional activation efficacy. In summary, we anticipate fluorescently labeled split-QF hemidriver systems will provide significant practical value and utility by enhancing reporter/effector expression specificity and thereby enabling functional characterization of a host of currently enigmatic cell types and subtypes.

## RESULTS

### Comparing tagged to untagged QF and QF2 holodrivers

To investigate the effects of adding a fluorescent tag independent of the effects of splitting the driver, we first tested whether fusing fluorescent reporters to an intact QF or QF2 holodriver impacts QUAS transcriptional activity. A red fluorescent protein (RFP), mScarlet specifically, was fused to the amino-terminus of QF and QF2 following a nuclear localization sequence (NLS). This allows direct comparison between driver and responder levels and highlights any cells that have high YFP expression but lack the corresponding RFP (false positives) or vice versa (false negatives; Fig. 1A). To assess relative transactivation efficacy, tagged or untagged holodriver constructs were injected into embryos of a stable transgenic responder line that expresses membrane-tagged yellow fluorescent protein (YFP), TagYFP2, under control of the QUAS (hereafter QUAS:YFP)^23^. At 3 days post-fertilization (dpf), confocal microscopy was used to collect images of YFP and RFP expression in cells in the caudal fin of injected larvae (Fig. 1B-F). Cellpose based segmentation was used to define RFP-expressing nuclei, YFP-expressing cells membranes, and quantify relative levels of each reporter per cell (Fig. 1G). For comparisons across each tested set here and below, measured intensities of each construct were normalized with the average of the construct yielding brightest YFP levels set to 1.

**Figure 1:**
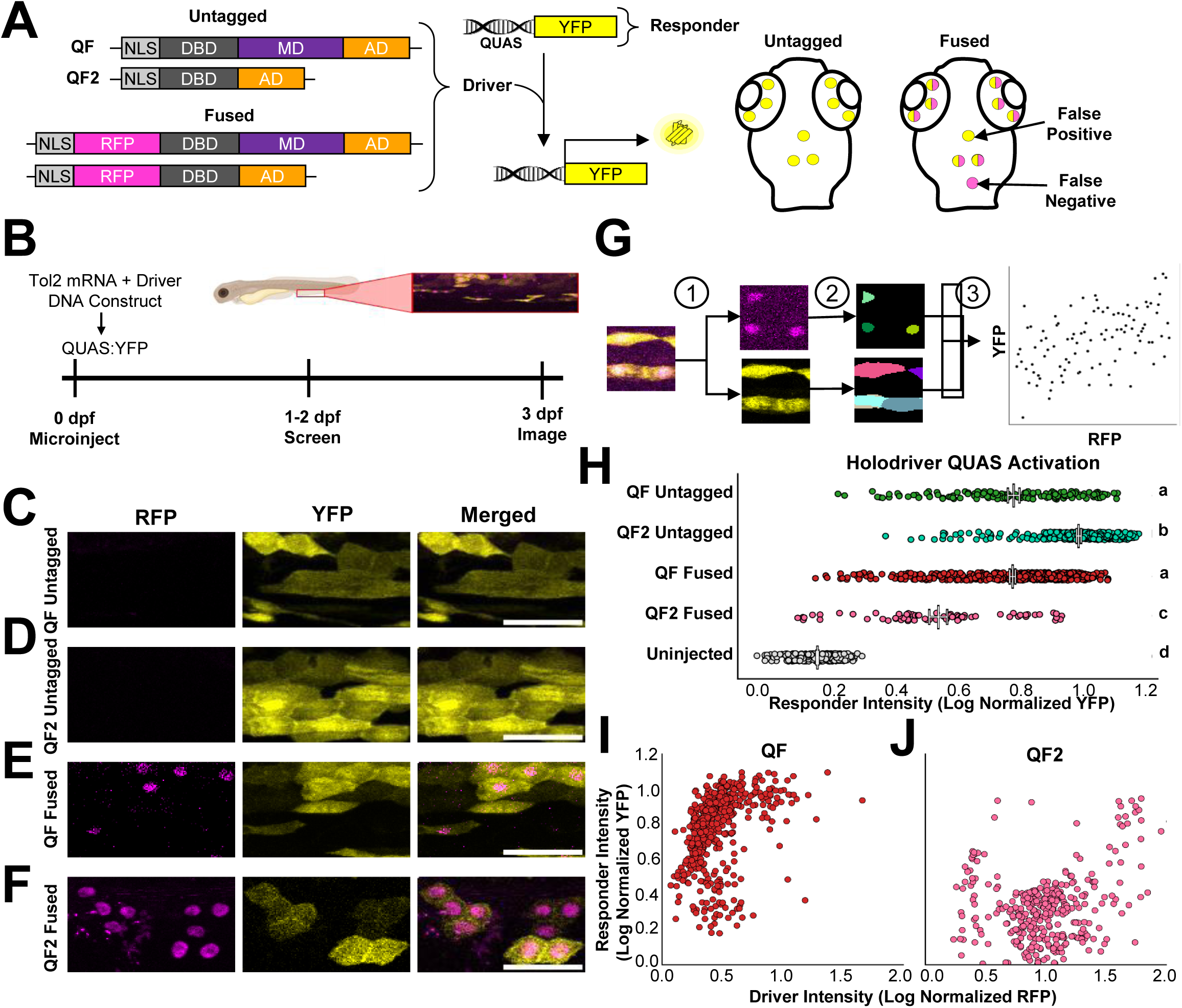
Mosaic genome integration paradigm to assess QUAS activation and evaluation of fluorescently tagged holodrivers. 1A) Schematic of untagged and tagged QF or QF2 “holodriver” DNA constructs that were used to drive YFP expression in QUAS:YFP larvae. While impossible with untagged holodrivers, RFP-tagged holodrivers identify false positives/negatives and facilitate direct comparison of driver and responder levels. DBD: DNA binding domain, MD: middle domain, AD: activation domain, NLS: nuclear localization sequence, YFP: yellow fluorescent protein, QUAS: QF upstream activating sequence. 1B) Workflow schematic of mosaic genome integration platform. Driver DNA constructs were injected with Tol2 mRNA into QUAS:YFP embryos at the single cell stage. The larvae were screened at 2 days post fertilization (dpf) for RFP and YFP expression. The brightest fish had a section of the caudal fin imaged on a confocal microscope at 3 dpf. 1C-F) Representative images of untagged and tagged QF and QF2 holodrivers injections, driving expression of the YFP in a mosaic manner. C,D; No RFP was observed when untagged constructs were injected. E,F; varying levels of nuclear RFP expression were observed in tagged construct injections. All constructs were able to cause YFP membrane expression in the epithelial cells of the caudal fin. Scale Bar = 50 µm. 1G) Schematic depicting fluorescence quantification pipeline. 1. Multicolor images into individual color channels, 2. Membranes and nuclei were segmented independently and 3. combined to define individual cells and corresponding nuclei, which the YFP and RFP intensity was extracted from. Individual cells were plotted as points on a scatter plot with RFP intensity on the X axis and YFP intensity on the Y axis. 1H) Strip chart comparing YFP intensity across tested groups as well as uninjected controls. Individual points represent average within independent ROIs from YFP membrane segmentation. Mean normalized YFP Intensity for Untagged QF Holodriver (0.80 ±0.018), untagged QF2 (1.00 ± .007), tagged QF (0.80 ± 0.008), and tagged QF2 (0.56 ± 0.028). Mean and SEM are depicted by white horizontal lines. a, b, c, and d denote statistical significance groups as identified by one way ANOVA with a Tukey’s Multiple Comparisons Correction. 1I-J) Scatterplot of QF(I) and QF2 (J) quantification with each cell as its own data point, demonstrating positive correlational relationship between RFP and YFP signal.

The untagged QF holodriver had lower mean normalized YFP intensity relative to the untagged QF2. Interestingly, the tagged QF2 holodriver showed reduced activity relative to its untagged control, whereas the tagged and untagged QF versions had no statistically significant difference in YFP expression (Fig. 1H). Additionally, tagged QF had stronger YFP expression than tagged QF2 across RFP levels, indicating that MD-containing holodrivers lead to increased responder activation when RFP is present (Fig. 1I,J). This data suggests that QF tolerates reporter tagging better than QF2. However, due to prior reports of QF driver toxicity when expressed broadly^17^, we elected to continue comparisons of QF and QF2 variants for hemidriver construct analyses.

### Hemidriver-reporter fusion constructs

To test the efficacy of hemidrivers compared to intact holodrivers, we injected untagged DBD-MD (QF) or DBD alone (QF2) constructs simultaneously with an AD (Fig. 2A,B). Images were collected and relative YFP levels were quantified and compared across groups (Fig. 2C). The QF hemidriver had slightly higher YFP levels in imaged fish than the corresponding untagged and tagged holodrivers, while the QF2 hemidriver had a markedly lower average YFP intensity than all three of these groups. These results support QF-based hemidrivers as a viable approach for reliable responder activation, and that QF is preferred over QF2 in the context of hemidrivers. However, given that this analysis was done in a G0 mosaic integration paradigm, it is impossible to determine whether differences in responder expression across groups was caused by differences in driver efficiency or variations in overall driver expression levels.

**Figure 2:**
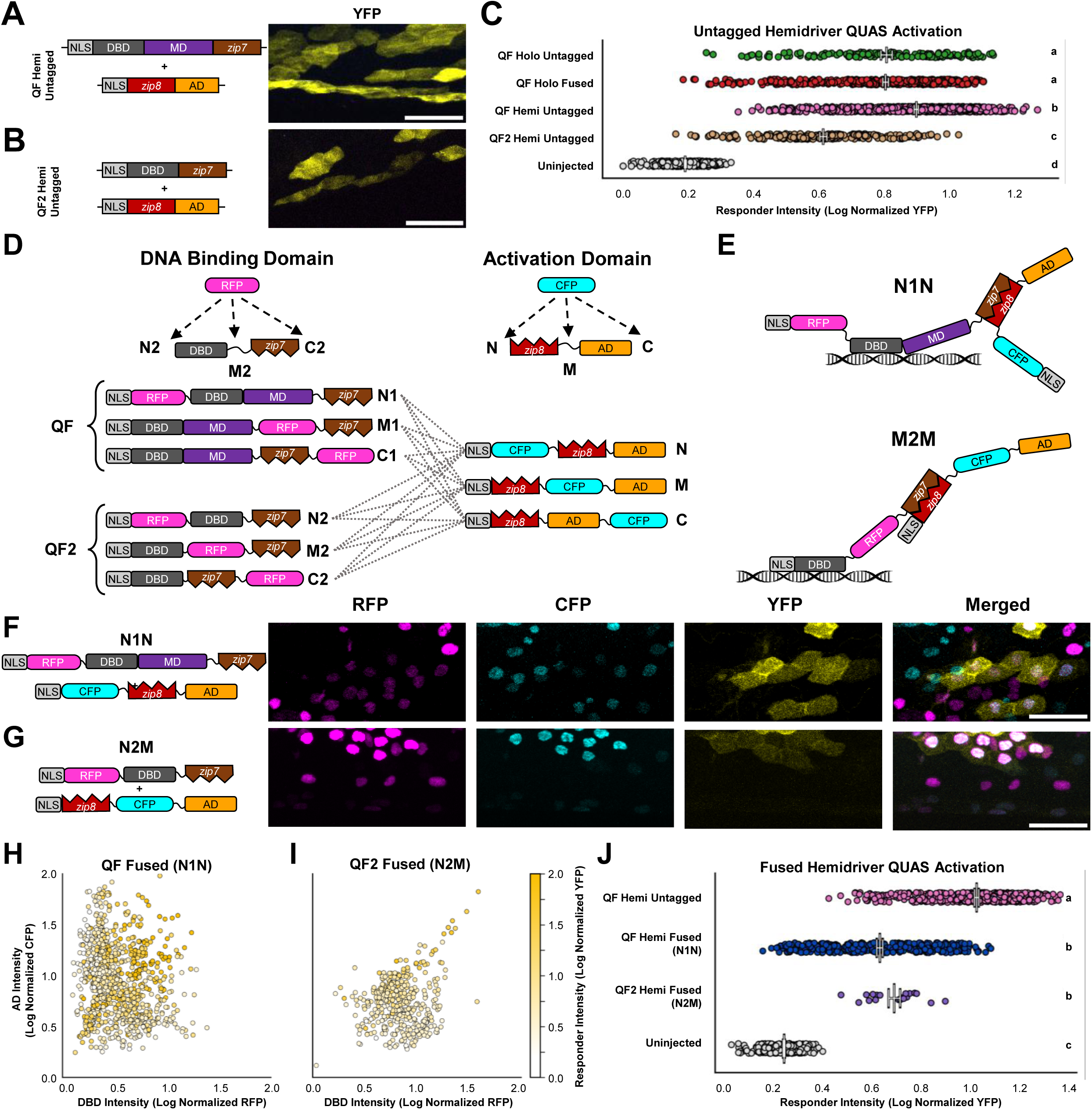
Hemidrivers and fluorescent tagging topology optimization. 2A-B) Schematic of untagged QF or QF2 hemidriver DNA constructs created by splitting holodrivers into DBD (with and without MD for QF and QF2 variations respectively) and AD components with a zip7 or zip8 dimerization domain added, respectively, and corresponding YFP channel images from injected larvae caudal fin. Scale Bar = 50 µm. 2C) Strip chart comparing YFP intensity from untagged hemidriver to results from tagged and untagged QF holodrivers (previous figure) as well as uninjected controls. As above, individual points represent average within independent ROIs from YFP membrane segmentation. Mean and SEM are depicted by white horizontal lines. a, b, c, and d denote statistical significance groups as identified by one way ANOVA with a Tukey’s Multiple Comparisons Correction. 2D) Schematic depicting possible topologies for tagged hemidriver protein constructs. A fluorescent protein is added in either the N-terminus, C-terminus, or between the dimerization domain and driver domains (termed M for Middle). mScarlet was attached to the DBD, while mTurquoise2 is attached to the AD. Constructs N1, M1, and C1 are QF DBDs that contain an MD, while constructs N2, M2, and C2 are QF2 DBDs that lack an MD. Each of the DBD constructs were tested in combination with each of the ADs, resulting in 18 total unique combinations. 2E) Schematic depicting complete driver formation when using tagged hemidrivers. Fluorophore placement impacts factors like total dimerized protein length, distance between DBD and AD, etc. 2F-G) QF and QF2 hemidriver combinations with the brightest YFP fluorescence (F, N1N for QF; G, N2M for QF2) protein schematic and representative confocal images showing each color channel individually and merged. Scale Bar = 50 µm. 2H-I) Three-dimensional scatterplots for the combinations shown in F and G. X and Y axes show normalized RFP and CFP intensity for segmented nuclei. YFP intensity from corresponding segmented membranes is represented as the hue of the point on a white to yellow color scale. 2J) Strip chart comparing YFP intensity from the brightest tagged QF and QF2 hemidriver to results from untagged QF hemidriver (Fig. 2A) as well as uninjected controls. As above, individual points represent average within independent ROIs from YFP membrane segmentation. Mean and SEM are depicted by white horizontal lines. a, b, c, and d denote statistical significance groups as identified by one way ANOVA with a Tukey’s Multiple Comparisons Correction.

To address this limitation in our untagged experiments, hemidrivers were fused to fluorescent reporters to support characterization of expression level in individual cells. The same RFP, mScarlet, was attached to DBD constructs, and a cyan fluorescent protein (CFP), mTurquoise, was used for AD constructs so that each hemidriver level could be quantified individually. To determine the optimal placement of the reporter in the hemidriver construct, variations were made in which the fluorescent protein was either at the N-terminus (N), between the DBD/AD and dimerization domains (“Middle”, M), or the C-terminus (C; Fig. 2D). For testing, QF-DBD variants (N1, M1, C1) and QF2-DBD variants (N2, M2, C2) were each co-injected with QF-AD hemidrivers (N, M, C), resulting in 18 unique combinations (e.g., N1N, M2M; Fig. 2E, see also Supp. File 1). Injections of individual constructs alone did not result in any YFP expression when images were collected at 3 dpf (Supp. Fig. 1), however co-injection of DBD and AD pairs led to robust YFP expression in the membrane of cells with corresponding hemidriver RFP and CFP expression (Fig. 2F,G).

The resulting images were analyzed as above, with the additional CFP channel combined with RFP for nuclear segmentation, and intensities quantified across all three channels (Supp. Fig. 2A). Three-way scatterplots were produced in which the X and Y axes show RFP (DBD) and CFP (AD) levels respectively, and the hue of the point on a white to yellow scale represents YFP intensity of that cell. This analysis showed that all pairs were able to activate the QUAS:YFP responder and confirmed positive correlations between DBD/AD co-expression levels and relative YFP levels (Fig. 2H,I; Supp. Fig. 3). When compared to untagged hemidrivers, even the combination with the brightest average YFP (N1N) was drastically lower than the untagged version (Fig. 2J), indicating further optimization would be necessary to produce a reliable tagged hemidriver.

Within a given cell, we are interested in YFP brightness relative to CFP/RFP brightness as a proxy for hemidriver expression levels rather than YFP brightness alone. To address this, we developed two additional analysis methods. In the first, an L-shaped bin defines the limiting hemidriver (CFP or RFP) and all points within this bin are averaged. The bin edges are then slid to higher limiting hemidriver levels, keeping the same width and allowing for overlap with the previous bin. This process is repeated until all possible values for RFP and CFP have been covered (Supp. Fig. 2C). The average YFP intensity across all points for each bin is then plotted against the bin center to produce a smooth curve that shows responder activation relative to the limiting amount of hemidriver (Supp. Fig. 2D). While this approach allows for comparison of YFP intensity across constructs for any given amount of hemidriver, the averaging has the potential to miss false positives or negatives in any group. To address this, a second approach was developed in which points were ranked into 10 groups (termed deciles) based on the level of limiting hemidriver expression (RFP or CFP). These deciles were allowed to vary in width but contained the same number of points (Supp. Fig. 2E). Each point within a given decile can then be plotted together with YFP intensity on the y axis, resulting in a graph where false positives and negatives become clear as deviations from the general trend of increasing YFP intensities in higher deciles (Supp. Fig. 2F).

The above analysis was run on all 18 combinations to determine the best fluorophore placement for a tagged hemidriver. The sliding bin line plots for QF (Fig. 3A) and QF2 (Fig. 3B) reveal clear frontrunners for each, N1N and N2M respectively, both of which have the highest average YFP intensities across all limiting hemidriver levels. The decile charts for these two combinations (Fig. 3C,D) reveal that lower deciles have averages slightly above background, but some bright YFP datapoints indicate strong QUAS activation even with limited amounts of hemidriver present. With higher deciles, the average YFP level increases as expected, but most importantly no points are observed at or below background (representing false negatives) in deciles 8-10. In contrast, two of the poorest performing combinations, M1C and C2M, points at or below background are observed in all 10 deciles (Fig. 3E,F). These decile plots were generated for all combinations (Supp. Fig. 4), and the first and last decile from each group was directly compared (Fig. 3G,H). Taken together, all analyses points towards N1N having the best topology for maximizing QUAS activation in a tagged hemidriver context, so we decided to move forward with this topology for further optimization.

**Figure 3:**
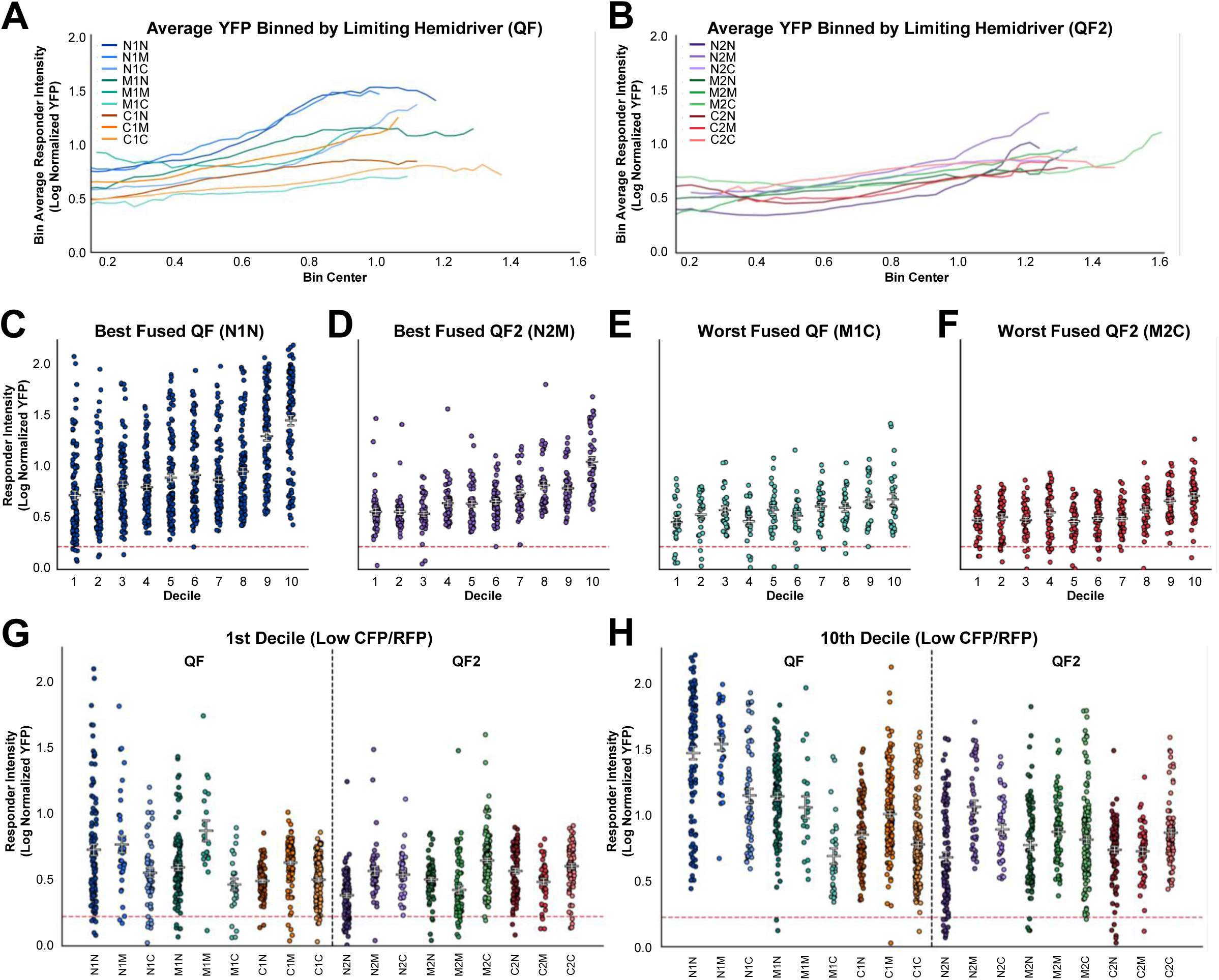
Assessment of fluorescent protein tagged topologies. 3A-B) Sliding window line plots depicting all QF (A) and QF2 (B) combinations tested. Graphs were produced using the approach outlined in supplementary figure 2C-D. X axis shows bin center, which represents the limiting amount of DBD or AD hemidriver present. Y axis shows average YFP intensity for all points within the given bin. 3C-F) Decile plots depicting most successful QF and QF2 combinations (C and D respectively) and least successful QF and QF2 combinations (E and F). Graphs were produced using the approach outlined in supplementary figure 2E-F. Horizontal red dashed line represents the average YFP intensity in uninjected controls. Mean and SEM for each decile depicted as white horizontal lines. 3G-H) Comparison of the first (G) and last (H) decile for all QF and QF2 combinations tested. Mean and SEM labeled per decile. Horizontal red dashed line represents the average YFP intensity in uninjected controls. Vertical black dashed line splits the QF and QF2 topologies.

### Shortened linkers enhance the transactivation efficacy of hemidriver-reporter fusions

Amongst our tagged hemidriver fusion constructs, we observed a general trend that the topologies which minimize the distance between the DBD and AD domains upon dimerization in the linear construct resulted in higher YFP expression. Therefore, we tested whether further minimizing this distance would result in increased YFP expression. The 10x poly-glycine linker previously connecting the DBD/AD and dimerization domains was shortened to a Gly-Ser linker in the top performing N1N hemidriver pair, denoted as N1N-Shortened Linker (N1N-SL; Fig. 4A). To account for general effects of the linker length and fluorescent tagging, we also created untagged hemidrivers with shortened linkers (untagged-SL) for comparison with the original linkers (untagged-OL). Hemidriver variants were injected in pairs, then imaged (Fig. 4B,C) and analyzed as above. In the untagged context, the SL variant had lower YFP levels on average, however, compared to the parent N1N construct, the N1N-SL showed increased YFP reporter intensity and followed the same proportional relationship between RFP/CFP and YFP intensity (Fig. 4D). When evaluating the line plots for these SL constructs, it becomes apparent that the YFP intensity for any given level of limiting hemidriver is relatively consistent, however the maximum amount of hemidriver measured was increased in both combinations tested, resulting in higher measured YFP (Fig. 4F). Additionally, the decile plot did reveal that the shortened linker version of N1N was more prone to false negatives, in which high levels of RFP and CFP were measured in the cell, but the YFP is at or below background levels (Fig. 4G). Together, these results suggest a complex relationship between hemidriver abundance and responder activation not explainable by length of the hemidriver alone. Overall, shortened linker improved tagged hemidriver-reporter expression through unknown mechanisms –perhaps improved folding kinetics or increased half-life– which, in turn, increased QUAS activation and ultimately YFP levels.

**Figure 4:**
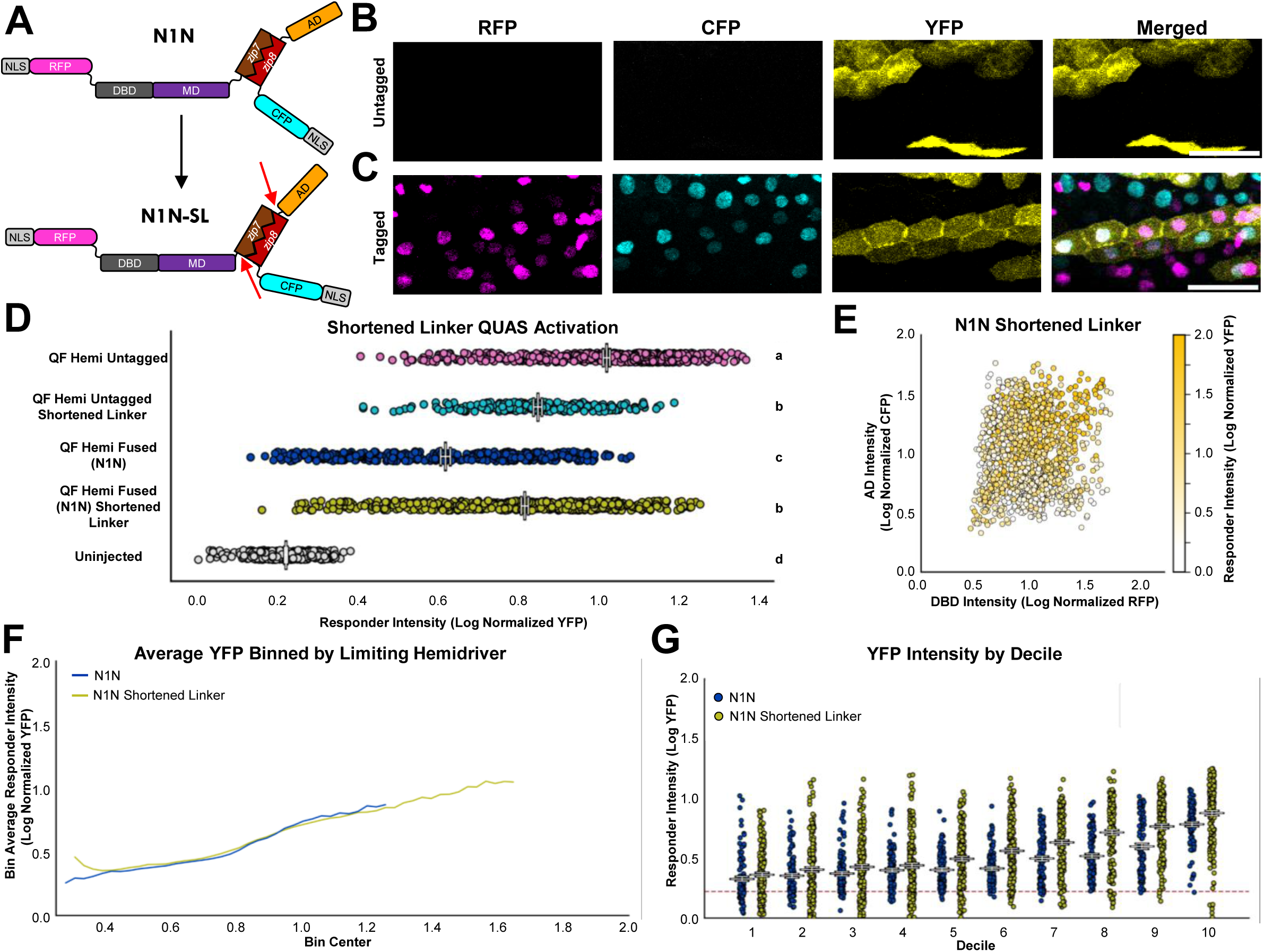
Fluorescent tagging optimization through linker sequence shortening. 4A) Schematic displaying the N1 and N proteins before and after linker sequence optimization. Linkers between the DD and DBD/AD were reduced in size. Red arrows indicate the regions that were shortened. 4B-C) Representative confocal images from caudal fin or larvae injected with untagged (B) and tagged (C) shortened linker variations of hemidriver constructs. Each color channel shown individually and merged. Scale Bar = 50 µm. 4D) Strip chart comparing YFP intensity from tagged and untagged QF hemidriver shortened linker constructs to results from using the original linker (Fig. 2A) as well as uninjected controls. As above, individual points represent average within independent ROIs from YFP membrane segmentation. Mean and SEM are depicted by white vertical lines. a, b, c, d, and e denote statistical significance groups as identified by one way ANOVA with a Tukey’s Multiple Comparisons Correction. 4E) Three-dimensional scatterplot for the N1N combination shown in C. X and Y axes show normalized RFP and CFP intensity for segmented nuclei. YFP intensity from corresponding segmented membranes is represented as the hue of the point on a white to yellow color scale. 4F) Sliding window plot showing the tagged shortened linker construct N1N-SL (yellow) as well as the data from the corresponding original linker, N1N (blue). 4G) Direct comparison of all deciles between the N1N shortened and original linker variations (yellow and blue, respectively). Horizontal red dashed line represents the average YFP intensity in uninjected controls. Mean and SEM for each decile depicted as white horizontal lines.

### Generation of hemidriver HaloTag fusion constructs

While linking hemidriver expression to fluorescent reporters is useful for the screening associated with the creation of transgenic lines and aids in assessing relative responder activation efficacy, the need for three complementary reporters crowds spectral space and limits compatibility with other reporter systems. To address this issue, we created hemidrivers linked to HaloTag, a monomeric enzyme derived from the bacterial gene *haloalkane dehalogenase* that covalently binds synthetic fluorescent dyes, to allow hemidriver-expressing cells to be labeled on an as-needed basis (e.g., to propagate transgenic lines or assess transcriptional activity fidelity). Both DBD and AD HaloTag conjugated constructs were produced and co-injected with the corresponding AD or DBD from our optimized N1N-SL pair from above. Injected fish were then incubated with Janelia Fluor HaloTag Ligand dyes spectrally distinct from the fluorescent protein tagged to the other hemidriver (Supp. Fig. 5A). Following washout of the dye fish were imaged and results were subject to the same analysis as above (Supp. Fig. 5B,C). Both versions yielded YFP averages slightly higher than the original version with both hemidrivers fluorescently tagged, yet not significantly different from each other (Supp. Fig. 5D). The HaloTag variants upheld the proportional relationship between hemidriver and responder levels (Supp. Fig 5E,F), and the sliding line plot reveals a nearly identical trend with the optimized N1N-SL combination (Supp. Fig. 5G). The decile plot shows an overall increased YFP expression in the HaloTag variants, especially in the middle deciles, indicating strong activation of the QUAS responder in both contexts (Supp. Fig 5H).

### Hemidrivers co-expressing fluorescent reporters using 2A peptides

Since QUAS:YFP activation efficacy was still relatively low in our optimized N1N-SL combination as compared to untagged hemidrivers, we sought to develop an approach to achieve a fluorescent hemidriver with activity closer to that of the untagged hemidriver. To avoid the potential for lower activity caused by reporter fusions, we tested the 2A peptide system for co-expressing fluorescent reporters with hemidrivers without physically linking them. In multi-cistronic 2A constructs, the expression level of some proteins has been shown to vary based on its position location within the construct^24^. We therefore elected to test hemidriver activity in both the N-terminal (Hemidriver-P2A-FP) and C-terminal (FP-P2A-Hemidriver) positions (Fig. 5A). DBD/AD construct variants did not cause any YFP expression when injected alone into the QUAS reporter line (Supp. Fig. 6), but all pairs tested led to robust YFP expression throughout the fish (Fig. 5B-E). As observed in the topology experiments, QF-P2A constructs had higher activity than QF2-P2A counterparts, and Hemidriver-P2A-FPs outperformed FP-P2A-Hemidrivers (Fig. 5F). Comparisons between P2A tagged construct variants showed strong positive correlations between CFP/RFP ratios and YFP intensities in all cases (Fig. 5G,H). Line and window plot analyses further indicated that the QF-P2A-FP variation produced the highest YFP intensity, leading us to deem this combo as the most efficient in terms of YFP expression relative to hemidriver levels (Fig. 5I,J).

**Figure 5:**
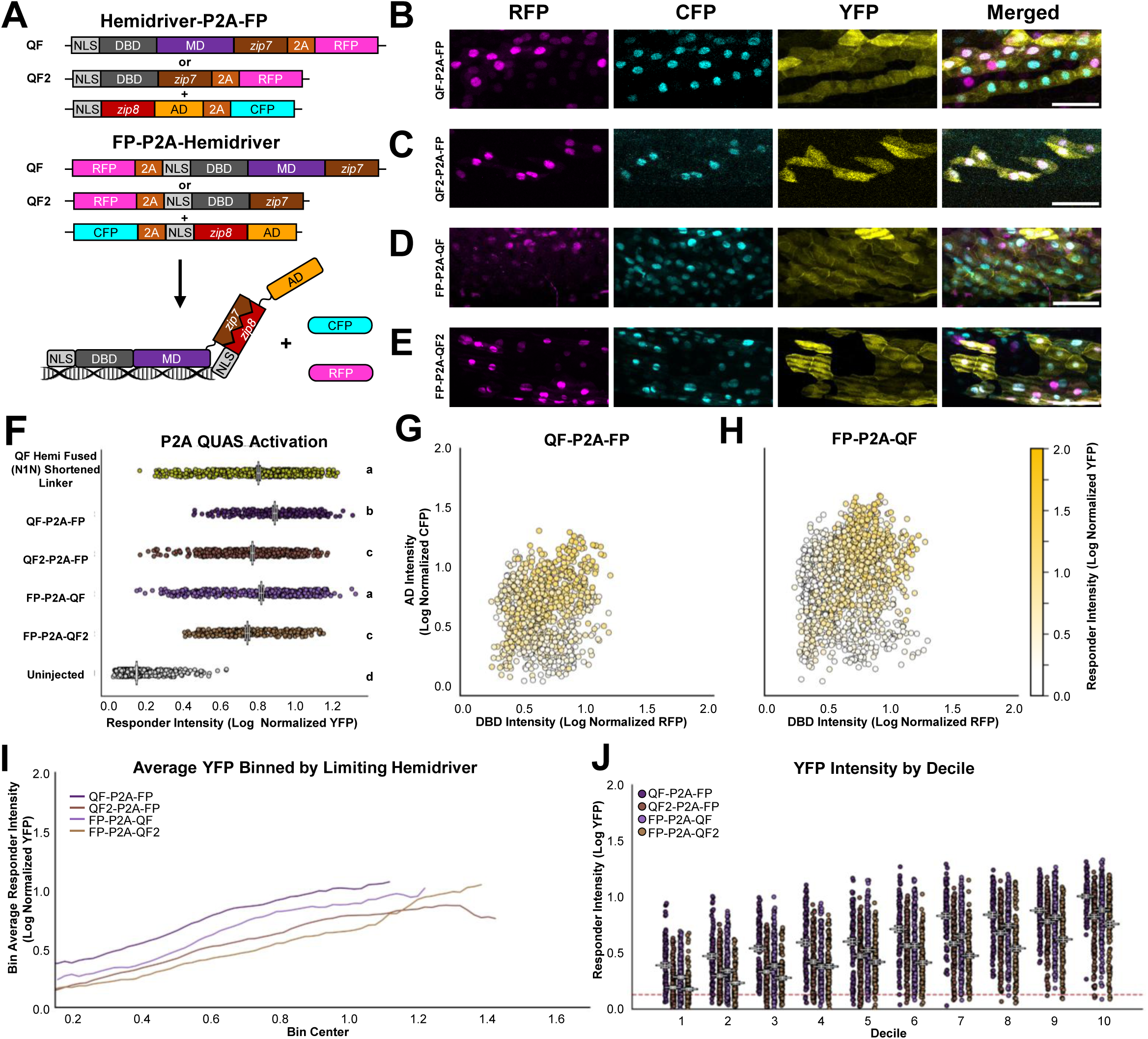
Hemidriver-fluorescent protein co-expression constructs using P2A peptides. 5A) Schematic of P2A peptide QF or QF2 hemidriver fluorescent protein co-expression DNA constructs created. Both C-terminal (Hemidriver-P2A-FP) and N-Terminal (FP-P2A-Hemidriver) variations were generated. Both result in free fluorescent protein that was cleaved from the corresponding hemidriver protein. 5B-E) Representative confocal images from the caudal fin of larvae injected with one of the four different combinations outlined in A. Each color channel shown individually and merged. Scale Bar = 50 µm. 5F) Strip chart comparing YFP intensity from P2A QF and QF2 hemidriver constructs to results from the optimized N1N-SL as well as uninjected controls. As above, individual points represent average within independent ROIs from YFP membrane segmentation. Mean and SEM are depicted by white vertical lines. a, b, c, d, e, and f denote statistical significance groups as identified by one way ANOVA with a Tukey’s Multiple Comparisons Correction. 5G-H) Three-dimensional scatterplots for the QF combination shown in F. X and Y axes show normalized RFP and CFP intensity for segmented nuclei. YFP intensity from corresponding segmented membranes is represented as the hue of the point on a white to yellow color scale. 5I) Sliding window plot showing all linker constructs tested (QF, purples, QF2, browns) as well as the optimized N1N-SL (yellow). 5J) Direct comparison of all deciles between P2A constructs and the N1N shortened linker tagged variation. Horizontal red dashed line represents the average YFP intensity in uninjected controls. Mean and SEM for each decile depicted as white horizontal lines.

### Structural analysis of QF holodrivers and hemidrivers

We identified several trends in activation efficacy that are evident across construct variants. First, in contrast to split-QF systems tested in *C. elegans* (which achieved only 42% activation relative to driver controls^25^), our top performing hemidrivers achieved similar levels of QUAS activation as their holodriver controls. Second, splitting the drivers had differing effects on the activation efficacy of QF and QF2. Although the untagged QF2 holodriver showed greater activation than the QF counterpart, untagged QF hemidrivers typically resulted in higher QUAS activation than their QF2 counterparts. Interestingly, the untagged QF hemidriver had increased activation compared to their holodriver counterpart while QF2 had decreased activation, suggesting that QF may better tolerate being split. Third, reporter fusions had minimal-to-no effect on QF variants but significantly reduced activation efficacy of QF2 variants for both holodrivers and hemidrivers, suggesting that QF constructs better tolerated FP-tags in zebrafish. Fourth, N-terminal tagged variants generally exhibited higher QUAS activation capacities, with N1N performing the best among hemidriver fusions. However, there were exceptions, such as N2M performing the best among QF2 variants. Lastly, shortening the linker sequence between the QF domain and zip domain in tagged hemidriver fusion variants increased their activation efficacy, but reduced activation of untagged hemidriver constructs. These trends, and the exceptions to them, highlight the unpredictability of construct design and the need for validating the efficacy of each hemidriver variant.

Given this unpredictability and the unknown nature of synthetic protein design, we next assessed if a structural analysis of different hemidrivers could provide further insight. Using AlphaFold 3, we first looked at how untagged and tagged QF holodrivers dimerize around a 1xQUAS sequence (Fig. 6A,B). In both cases, we can see the DBD successfully binding to the QUAS. In the tagged version, we noted that the FP is situated opposite the MD, balancing the overall structure around the QUAS. With this understanding, we next explored the structures of the best and worst QF hemidrivers, N1N and M1C respectively. We found that the majority of domains had low expected error in their predictions (FPs, DBD, MD, and the Zip Domains), while the AD consistently displayed a highly disordered region with a higher expected position error (Fig. 6C), consistent with prior descriptions of this domain^26^. Akin to the QF tagged holodriver (Fig. 6B), N1N adopted a similar overall structure with the FPs being situated opposite the MD (Fig. 6D), while M1C showed the FPs placed next to the MD, disrupting the balanced domain topology observed for N1N and tagged driver controls and potentially creating steric hinderance (Fig. 6E). Nevertheless, despite M1C seeming to have a larger gap between the two dimers, these changes did not seem to impact DBD binding to the QUAS (Fig. 6F,G), suggesting that steric hinderance of the DBD is not a main driver of the differences in QUAS activation between topologies. Additionally, the zip domains were able to come together and showed similar hydrogen bonding patterns in both topologies (Fig. 6H,I), suggesting that variability in the AD- and DBD-hemidrivers coming together is also not a main driver of transactivation differences. Given these results, along with the trend that hemidriver pairs with C-terminally tagged ADs showed lower QUAS activation efficacy, we speculate that differences in activation efficiency are likely due to variations in the AD’s ability to recruit transcriptional machinery to the QUAS, potentially due to steric hinderance resulting from FP placement within the overall protein structure.

**Figure 6:**
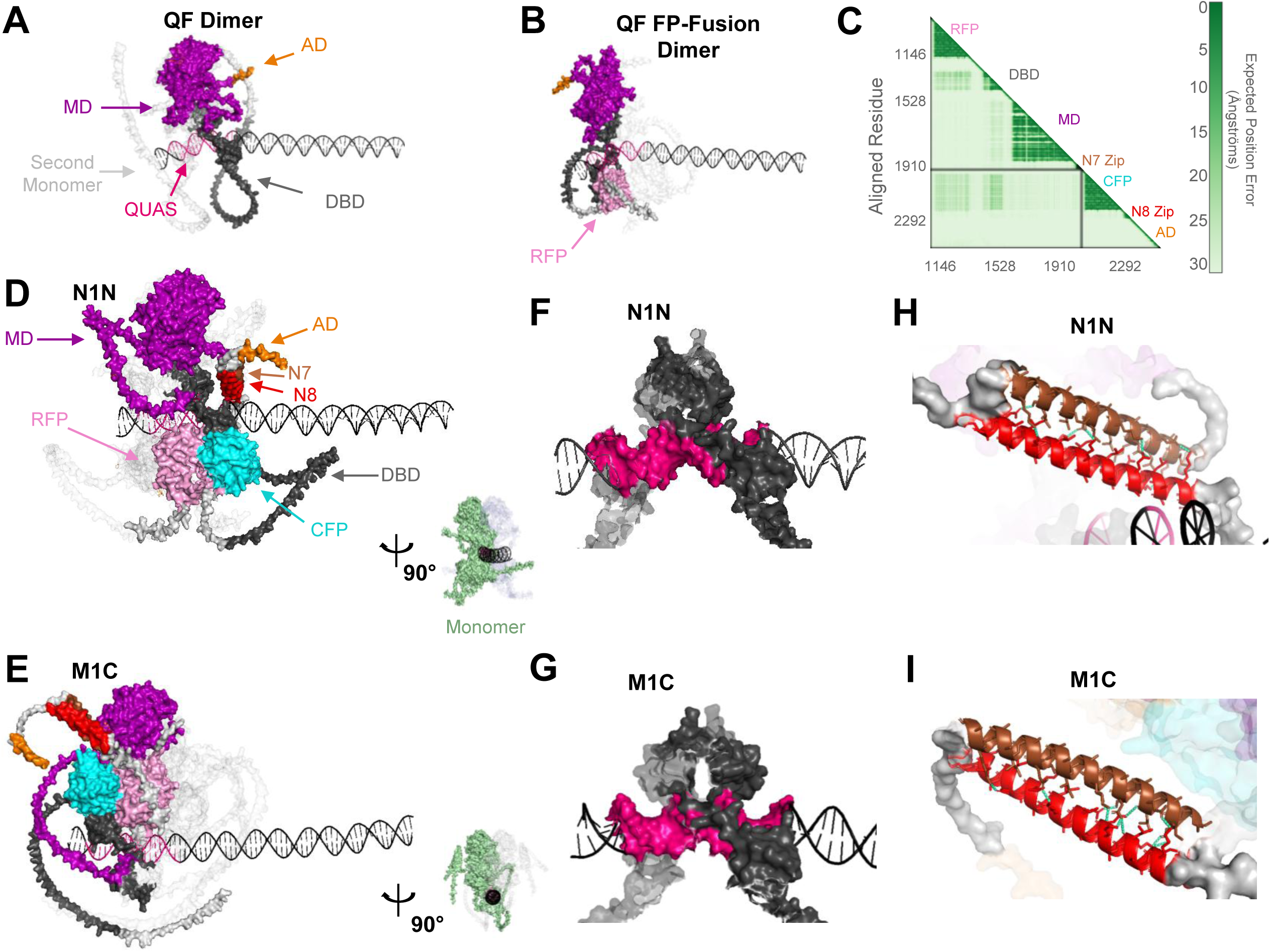
Structural Analysis of QF Holodrivers and Hemidrivers. 6A-B) Structural prediction of the untagged QF driver as a dimer (A) and QF driver FP-Fusion as a dimer (B) binding to a 1x QUAS DNA sequence. 6C) Scored Residues of N1N with domains labeled. 6D-E) Topology of a top performing hemidriver, N1N (D), and a bottom performing hemidriver, M1C (E), with domains labeled, with 90° rotated view of N1N and M1C looking down the DNA helix, showing one monomer in green. 6F-G) The DBD binding of N1N (F) and M1C (G) binding the QUAS in the minor groove. 6H-I) Hydrogen bonding, shown in green-cyan, between the N7 (brown) and N8 (red) zip domains in N1N (H) and M1C (I). For clarity, the disordered region of the AD Domain (orange) has been visually truncated, and one monomer is displayed transparently.

### Summary – comparison of best performing variants

For ease of comparison, we plotted all best performing split-QF constructs in a single strip chart, sliding line plot, and decile chart to correlate relative QUAS:YFP activation efficacy across variants (Fig. 7A-D). Maximal QUAS activation is achieved by both untagged hemidrivers and 2A co-expression hemidrivers, specifically the QF-2A-FP topology, while reporter fusions resulted in diminished activity that can be partially rescued by a shortened linker. While untagged hemidrivers achieve maximal QUAS activation levels, tagged hemidrivers provide the ability to trace the expression pattern of individual hemidrivers, which offers the advantages of (i) facilitating the establishment of novel transgenic lines, (ii) supporting *in vivo* visualization of multiple marker genes, (iii) detecting instances of failed/spurious activation, (iv) accounting for any expression changes that might arise over time, and (v) accounting for expression variation between individual fish per experiment, providing an important option for simplifying line creation and maintenance, and increasing resource and assay robustness.

**Figure 7:**
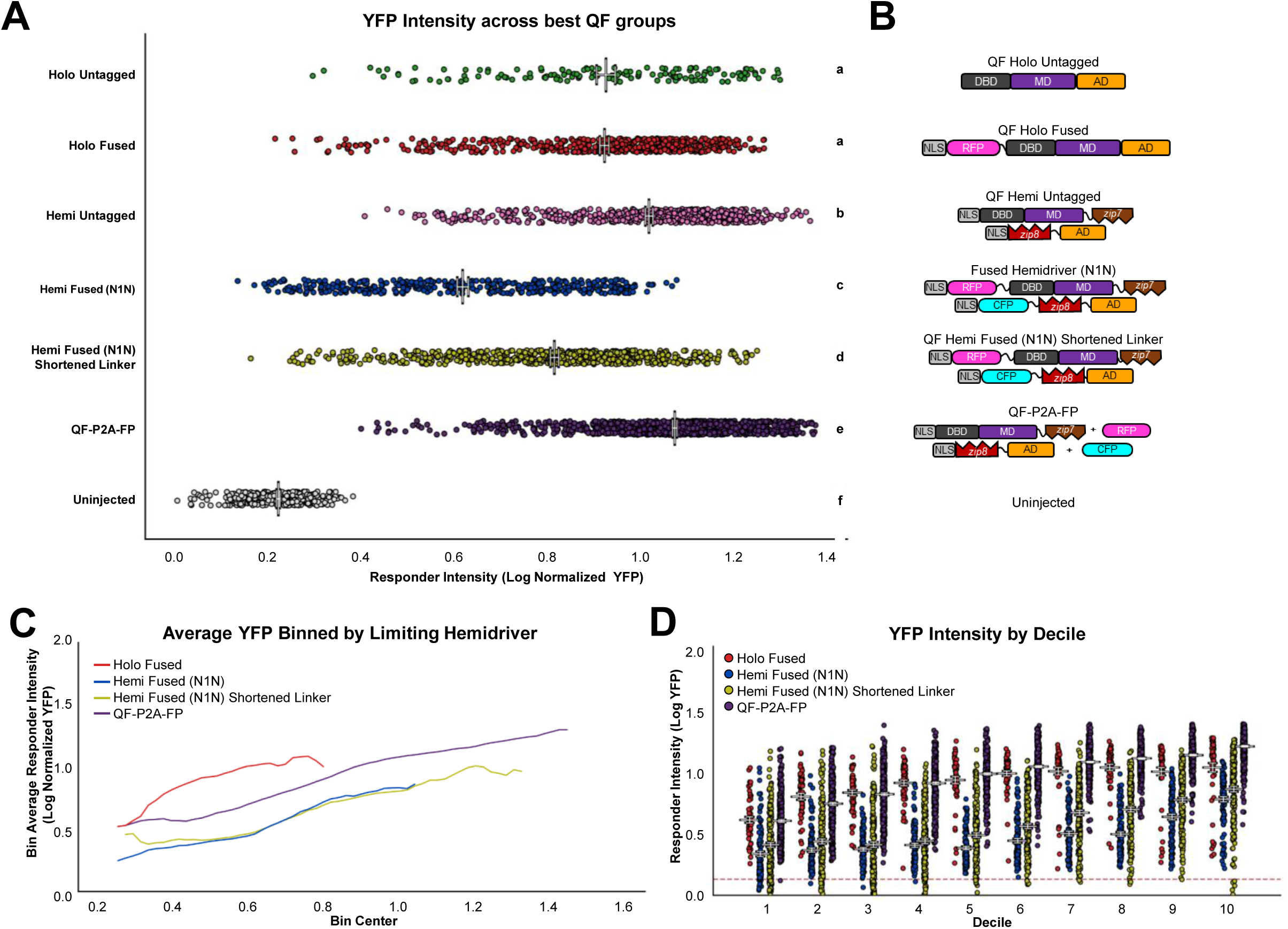
Summary Comparison of QUAS Activation Across QF Constructs. 7A) Strip chart comparing YFP intensity across the best QF construct from each group of holo/hemidrivers. As above, individual points represent average within independent ROIs from YFP membrane segmentation. Mean and SEM are depicted by white vertical lines. a, b, c, d, e, and f denote statistical significance groups as identified by one way ANOVA with a Tukey’s Multiple Comparisons Correction. 7B) Schematic of each QF driver/hemidriver construct depicted in 6A. 7C) Sliding window plot showing the average YFP intensity all tagged constructs depicted in 6A, namely the QF fused holodriver (red), QF fused hemidriver N1N (blue), shortened linker QF fused hemidriver N1N-SL (yellow), and QF-P2A-FP (purple). 7D) Direct comparison of all deciles of YFP intensity between the tagged constructs depicted in 6A, namely the QF fused holodriver (red), QF fused hemidriver N1N (blue), shortened linker QF fused hemidriver N1N-SL (yellow), and QF-P2A-FP (purple).

## DISCUSSION

Split-driver systems enable improved reporter/effector transgene expression specificity through intersectional targeting of discrete cell types. The advent of single cell transcriptomics provides both a challenge and an opportunity in this regard, increasing the number of predicted cell types but also providing genetic combinations for selectively targeting potential “new” cells. Combined with efficient knock-in methods, intersectional strategies can be used to generate stable transgenic lines facilitating exclusive labeling and manipulation of transcriptomics-predicted cell types. In turn, these resources enable functional tests to resolve predicted cell type specific roles and thereby refine cell type definitions. As examples, extensive split-driver toolsets, derived largely by enhancer trap selection, have been used to reveal core principles of neural circuit design and function by enabling exclusive targeting of discrete neuronal subtypes in flies and worms^9,27–33^. Using an intersectional targeting approach combining QF driver lines, a Cre recombinase line, and Cre-dependent QUAS reporter/effectors, Kolsch et al., assessed neuronal subtype function in the zebrafish retina^1^. Combining split-drivers with recombinase-based systems expands intersectional targeting options by enabling 3- and 4-way targeting. This is predicted to allow an additional 41% of single cell transcriptomics-defined neuronal cell types to be selectively targeted – adding ∼34% by 3-way and ∼7% by 4-way logic gates; in sum, >98% coverage of all predicted cell types across single, paired, ternary, and quaternary targeting options^22^. Achieving 3-way targeting will be particularly important for adequately targeting functionally distinct neuronal cell subtypes. Recent studies suggest a third spatial targeting element will be required to account for observations that binary genetic combinations are often shared between neuronal types exhibiting functional diversity (i.e., differential responses to stimuli)^20,22^ but which resolve to distinct functional profiles when looking at responses in a specific brain region (e.g., somal lamina). Accordingly, adding split-drivers to vertebrate molecular biology toolboxes will expand intersectional targeting options, supporting next-level transgene expression specificity enabling functional dissections of potential novel cell types revealed by single cell transcriptomics.

Although a split-Gal4 system was shown to be operable in zebrafish larvae over ten years ago^13^, the Kolsch et al. report is the sole example of functional cell type analysis using intersectional targeting in zebrafish. Similarly, though dual recombinase systems are used to achieve higher-order transgene expression specificity in mice and rats, split-drivers have yet to be incorporated into mammalian transgenic toolsets. Several practical barriers limit widespread adoption of split-driver methods, perhaps the most significant being the requirement to derive and maintain split-driver lines in ternary combinations of AD, DBD, and UAS/QUAS components in order to express fluorescent reporters for visual selection. To address this barrier and enhance intersectional targeting capabilities for vertebrate models, we developed novel fluorescently labeled split-driver resources. We chose to use the QF/QUAS system^8^ due to the potential to reduce transgenerational epigenetic silencing issues evident in UAS lines^14,15^, and because split-QF drivers had yet to be vetted for vertebrate systems. Reporter labeling involved either hemidriver-reporter fusions or 2A viral peptides for co-expression. Labeling split-driver components allows AD and DBD lines to be generated and maintained independently, thus enabling unique AD-DBD combinations to be brought together more efficiently. Tagging also accounts for any expression changes that might arise over time and, more importantly, expression variation between individual fish per experiment, providing an important internal control that increases intersectional assay robustness. Using complementary reporters (e.g., CFP and RFP variants) allows AD-DBD targeting to be visualized directly rather than computationally estimated. This also enables activation efficiency and off-target activity to be accounted for during resource optimization, as exemplified here in comparisons of 18 topological construct variants, and over the life of transgenic lines. Finally, we explored the use of a dye-labeled reporter option (e.g., HaloTag) to allow broad control of reporter “color” choice and enhance utility. These chemigenetic systems allow quality control assays (activation fidelity and targeting visualizations) using dye labels to be performed separately, circumventing the need to label hemidrivers during experimental assays. The full range of color channels used for *in vivo* imaging is thereby available for combining the power of intersectional targeting precision with existing transgenic resources. Additional factors must be considered when selecting among labeled split-driver options for a specific purpose. Accordingly, we recommend selecting hemidriver variants depending on the precise application (for examples, see Supp. File 14).

In summary, split-driver systems enable improved “reporter/effector” transgene expression specificity via two-way logic gates. Combined with other intersectional systems, such as recombinases (Cre, Cre^ERT2^, Flp etc.) or elements that can inhibit driver activity (e.g., QS), greater spatial and temporal control of transgene expression can be achieved by three- and four-way logic gates, thereby enabling targeting of nearly all potential novel cell types revealed by single cell transcriptomics The labeled split-driver resources presented here address practical bottlenecks hindering widespread adaptation, and better account for experimental variability, increasing robustness. Expanding the adaptability and utility of split-driver toolsets to vertebrate model systems will advance understanding of novel cell type function and interactions between discrete cell types, such as the logic of neural circuit design and function in vertebrate brains.

## MATERIALS AND METHODS

### Ethics Statement

Zebrafish protocol was approved by the Animal Care and Use Committees of the Johns Hopkins University School of Medicine. All zebrafish experiments were conducted in accordance with both the Association for Research in Vision and Ophthalmology (ARVO) statement on the “Use of Animals in Ophthalmic and Vision Research” and the National Institutes of Health (NIH) Office of Laboratory Animal Welfare (OLAW) policies regarding studies conducted in vertebrate species.

### Zebrafish maintenance

Zebrafish were maintained at 28.5°C with a light cycle of 14 hours of light and 10 hours of dark. Injections were done into a transgenic line, *Tg(5xQUAS:GAP-tagYFP-2A-nfsB_Vv F70A/F108Y,he:tagECFP)jh562* (hereafter QUAS:YFP) which expresses a membrane tagged yellow fluorescent protein (YFP)^23^. When combined with the QF hemidriver injections to drive expression of the QUAS, the YFP allows for the visualization and quantification of neurons cells. This line was outcrossed to a pigmentation mutant with reduced iridophore numbers, *roy^a^*^9^ (*roy*), and treated with 1 × PTU (TCI Chemicals, P0237) to facilitate YFP reporter signal detection *in vivo*.

### Construct Generation

DNA fragments with BsaI extensions were PCR-amplified from existing plasmid backbones or gene syntheses using Phusion High-Fidelity DNA Polymerase (New England Biolabs, M0530). After size validation by gel electrophoresis, fragments were gel extracted (Zymo Research, D4008) and ligated together using BsaI Golden Gate assembly (New England Biolabs, E1602). Constructs were transformed into competent DH5α E Coli cells (New England Biolabs, C2987H), then purified via mini-prep (Macherey-Nagel, 740588) and validated using whole plasmid Nanopore sequencing (Quintara Biosciences). All constructs generated and used in this study can be found in Supp File 2.

### Tol2 Injections and Screening

A mixture of hemidriver component plasmids (5 ng/µL each), Tol2 transposase mRNA (10 ng/µL; See Fig S1B), and phenol red (Sigma-Aldrich, P3532) were injected into single cell embryos. At 2 days post fertilization (dpf), the injected larvae were screened for a QUAS-CFP-hatching enzyme tracer, QUAS-YFP signal, and hemidriver RFP and CFP signal. Larvae with YFP signal, or overlapping RFP and CFP signal without YFP signal, were selected for confocal imaging and fluorescence quantification.

### Imaging

Fish were anesthetized in 1x Tricaine Methanesulfonate (Syndel, MS-222) and then embedded laterally in 1% low-melting agarose (Fisher BioReagents, BP1360-100) diluted in E3 and imaged with on either an Olympus FV1000 confocal microscope (upright), equipped with a XLUMPlanFl 20×/0.95 water immersion objective (Figures 1-4,6) or an Olympus FV4000 confocal microscope (inverted), equipped with a UPlanSApo 30×/1.05 silicon oil immersion objective (Figure 5). Three-channel confocal z-stacks centered on the ventral fin were collected with a 5-µm step-size. Each channel was imaged individually in a sequential manner to minimize bleed through. See table below for excitation/emission specifics.

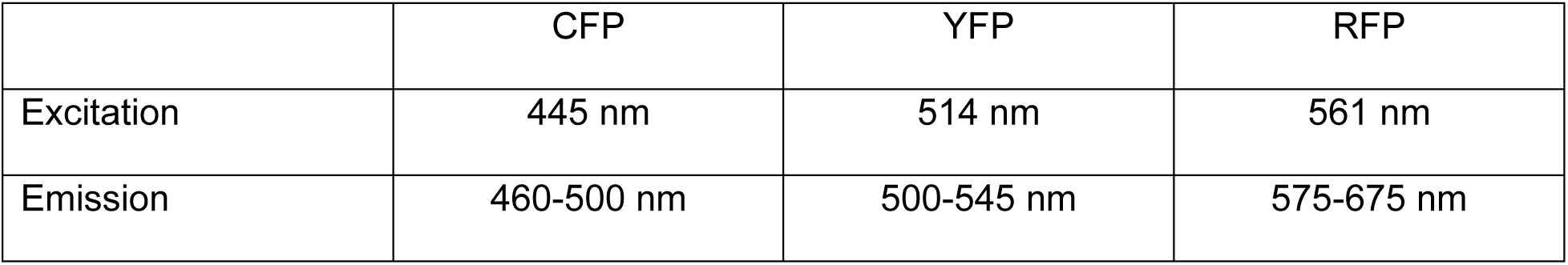

### Nuclei Segmentation and Fluorescence Quantification

First, a custom python preprocessing script split images into individual color channels and recombined the two nuclear channels based on maximum intensity per pixel following normalization to account for differences between CFP and RFP channels. Segmentation was conducted on these single channel z-stacks using CellPose^34^, with separate models for nuclei and cell membranes, each self-trained using ∼10 manually annotated single slices from the combined CFP/RFP channel or YFP channel respectively. Slices were segmented individually and combined in 3D based on a minimum overlap threshold using the parameters do_3D=False and stitch_threshold=0.2. The resulting segmentation masks were used for subsequent quantification using custom python scripts. ROIs defined by segmentation were classified as matched (segmented membrane with corresponding segmented nuclei within), orphan nuclei (segmented nuclei with no surrounding membrane), or orphan membrane (segmented membrane with no nuclei within). Intensities were extracted from the raw input image following global background subtraction and averaged over the entire region of interest defined by segmentation masks. For matched cells, CFP/RFP intensities were extracted from the region defined in the nuclei mask, and YFP intensity was extracted from the region defined by the corresponding membrane. For orphan nuclei and membranes, a single ROI from either the nuclear or membrane mask was used for extracting intensity for all three channels. These three intensity values for a given ROI as well as its segmentation status (matched, orphan cell/nuclei) were stored in CSV format for subsequent analysis.

### Data Visualization and Statistical Analysis

Custom Jupyter Notebooks were developed for analysis of all data. Logarithmic transformation was first applied to input intensities to create a normal distribution. For each set of comparisons, all channels were normalized, such that the average of the brightest group was set to 1 for each channel. Plots were generated using Seaborn python package^35^. In plots only showing YFP intensity, ROIs characterized as orphan nuclei were omitted. One-way ANOVA was performed on these YFP intensity graphs using scipy.stats.f_oneway and Tukey’s Honest Significant Difference test was implemented using statsmodels.stats.multicomp.pairwise_tukeyhsd with an alpha value of 0.05.

For sliding window line plots, a bin width of defined by dividing the range in values by 8 (approximately 0.25 for each bin plot)was used with an overlap of 0.9 between neighboring bins, and upper and lower limits for all bins were defined that stretched from the minimum to maximum value for CFP/RFP in the observed data. For each bin, the YFP intensity of all points within was averaged and plotted against that bin center; bins with less than 5 points were omitted. For decile charts, a new column was created that contained the minimum between the CFP and RFP columns. Data was then reordered based on this column, and pd.qcut (q=10) was used to divide the resulting dataframe into 10 equal groups based on this column.

### AlphaFold 3 and PyMOL Structural Analysis

QF driver and hemidriver amino acid sequences (See Supp File 2) and a 1xQUAS sequence

“GTCGACGGATAAACAATTATCCTCGCAAGGGTCGACTCTAGAGGGTATATAATGGATCCCATCGCGTCT”, and its compliment were inputted into AlphaFold 3^36^ as homodimers. The resulting AlphaFold3-generated PDB files were downloaded, and the model_0 versions were used for PyMOL analysis. Scripts for each construct were written and run to label each monomer, domain, and interaction and can be found in Supp Files 10-13.

## DATA AVAILABILITY

All plasmids generated have been submitted to Addgene. The code used for analysis is available on the mummlab github (https://github.com/mummlab/Hemidriver). All source code and raw images are available in the supplemental files. All AlphaFold 3 Structures have been uploaded to ModelArchive (https://modelarchive.org/) and PyMOL scripts can be found in the Supp Files 10-13.

## SUPPLEMENTAL FIGURE LEGENDS

**Supplemental Figure 1: Individual tagged hemidriver solo injections**

Representative images from negative controls in which either DBD or AD were injected by themselves. YFP channel shown from all images to show that no hemidriver alone caused YFP expression. Corresponding three-dimensional scatterplots showing segmented nuclei are depicted to the right.

**Supplemental Figure 2: Multicolor image analysis methods**

S2A) Confocal images are separated into component color channels, nuclei and membrane channels segmented separately then combined, and a single nucleus is assigned to a single segmented membrane.

S2B) Three-dimensional scatterplot to depict segmentation results. X and Y axes are RFP and CFP intensity respectively. Yellow intensity represented by hue of point.

S2C) Sliding window analysis performed on three-dimensional scatterplots. Define standard bin size for grouping based on range of hemidriver fluorescent protein intensities. Slide bin to higher CFP/RFP intensities until all points are covered

S2D) Plot of average YFP of all ROIs within a given bin versus bin center.

S2E) Decile analysis performed on three-dimensional scatterplots. Sort ROIs based on limiting hemidriver. Divide into 10 non-overlapping “deciles” with equal number of points.

S2F) Plot average YFP intensity for each ROI within every decile.

**Supplemental Figure 3: Tagged hemidriver topology experiments all three-dimensional scatterplots**

Three-dimensional scatterplots for all tagged fusion hemidriver topologies. X and Y axes show normalized RFP and CFP intensity for segmented nuclei. YFP intensity from corresponding segmented membranes is represented as the hue of the point on a white to yellow color scale.

**Supplemental Figure 4: Tagged hemidriver topology experiments all decile plots**

Comparison of all deciles for all tagged fusion hemidriver topologies. Horizontal red dashed line represents the average YFP intensity in uninjected controls. Mean and SEM for each decile depicted as white horizontal lines.

**Supplemental Figure 5: Replacement of fluorescent protein tag on hemidriver with HaloTag**

S5A) Schematic of DBD-HaloTag and AD-HaloTag hemidriver variants in combination with the AD N-SL and DBD N1-SL respectively. HaloTag ligands are depicted by stars attached to the HaloTag in each construct.

S5B-C) Representative confocal images from caudal fin or larvae injected with DBD-HaloTag + N-SL (B) and AD-HaloTag + N1-SL (C) and stained with a HaloTag ligand, JF-522 (B) or JF-647 (C). Each color channel shown individually and merged. Scale Bar = 50 µm.

S5D) Strip chart comparing YFP intensity from DBD-HaloTag and AD-HaloTag hemidriver constructs, in combination with N-SL and N1-SL respectively, to results from the N1N-SL constructs (Fig. 4A) as well as uninjected controls. As above, individual points represent average within independent ROIs from YFP membrane segmentation. Mean and SEM are depicted by white vertical lines. a, b, c, and d denote statistical significance groups as identified by one way ANOVA with a Tukey’s Multiple Comparisons Correction.

S5E-F) Three-dimensional scatterplots for the HaloTag combinations shown in A. X and Y axes show normalized RFP and CFP intensity for segmented nuclei. YFP intensity from corresponding segmented membranes is represented as the hue of the point on a white to yellow color scale.

S5G) Sliding window plot showing the HaloTag construct combinations tested (DBD-HT, red, AD-HT, orange) as well as the optimized N1N-SL (yellow).

S5H) Direct comparison of all deciles between HaloTag construct combinations and the N1N shortened linker tagged variation. Horizontal red dashed line represents the average YFP intensity in uninjected controls. Mean and SEM for each decile depicted as white horizontal lines.

**Supplemental Figure 6: Individual P2A hemidriver solo injections**

Representative images from negative controls in which either a P2A DBD or AD were injected by themselves. YFP channel shown from all images to show that no hemidriver alone caused YFP expression. Corresponding three-dimensional scatterplots showing segmented nuclei are depicted to the right.

## ACKNOWLEDGEMENTS

The authors thank the Wilmer Core facilities for their important contributions to this work, as well as the unrestricted departmental grants to the Wilmer Eye Institute. The authors additionally thank the members of the Mumm and Thierer labs for helpful discussion. Some schematics for figures were created with Biorender.com.

