## Supplemental Figures for "Fluorescently-labeled split-QF hemidrivers: simplifying and enhancing methods enabling intersectional targeting of discrete cell types"

### Supplemental Figure 1: Individual tagged hemidriver solo injections

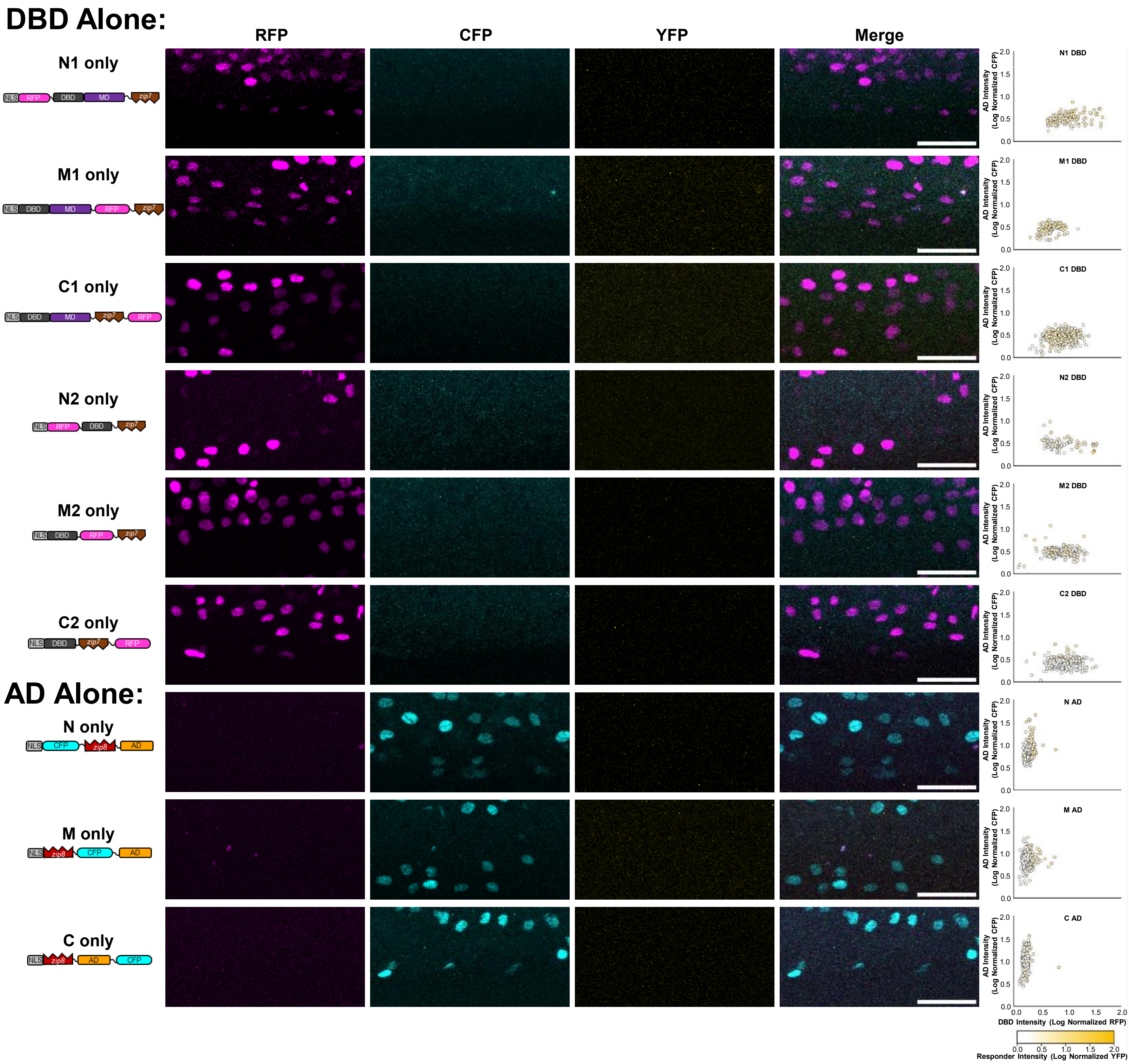

### Supplemental Figure 2: Multicolor image analysis methods

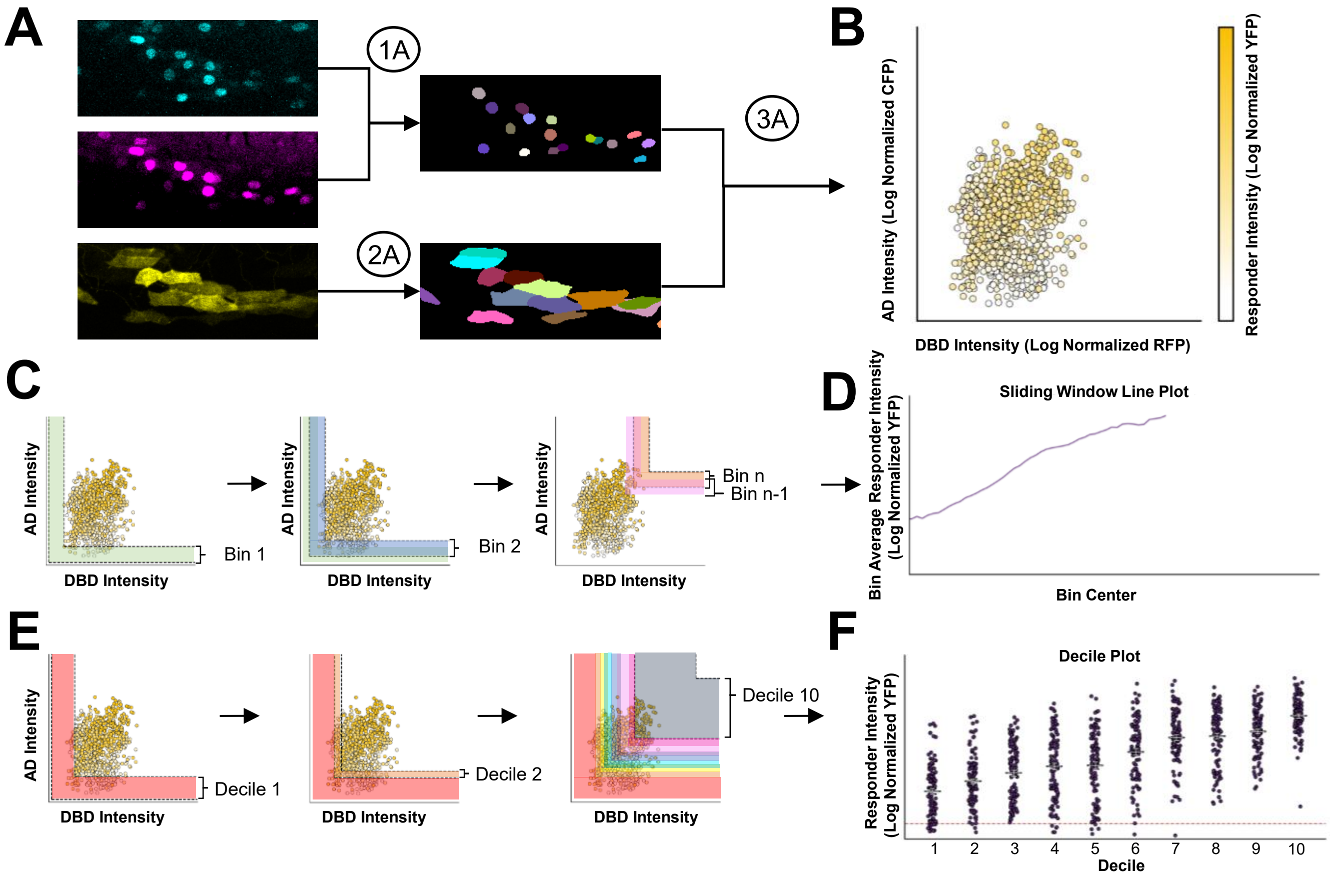

### Supplemental Figure 3: Tagged hemidriver topology experiments all three-dimensional scatterplots

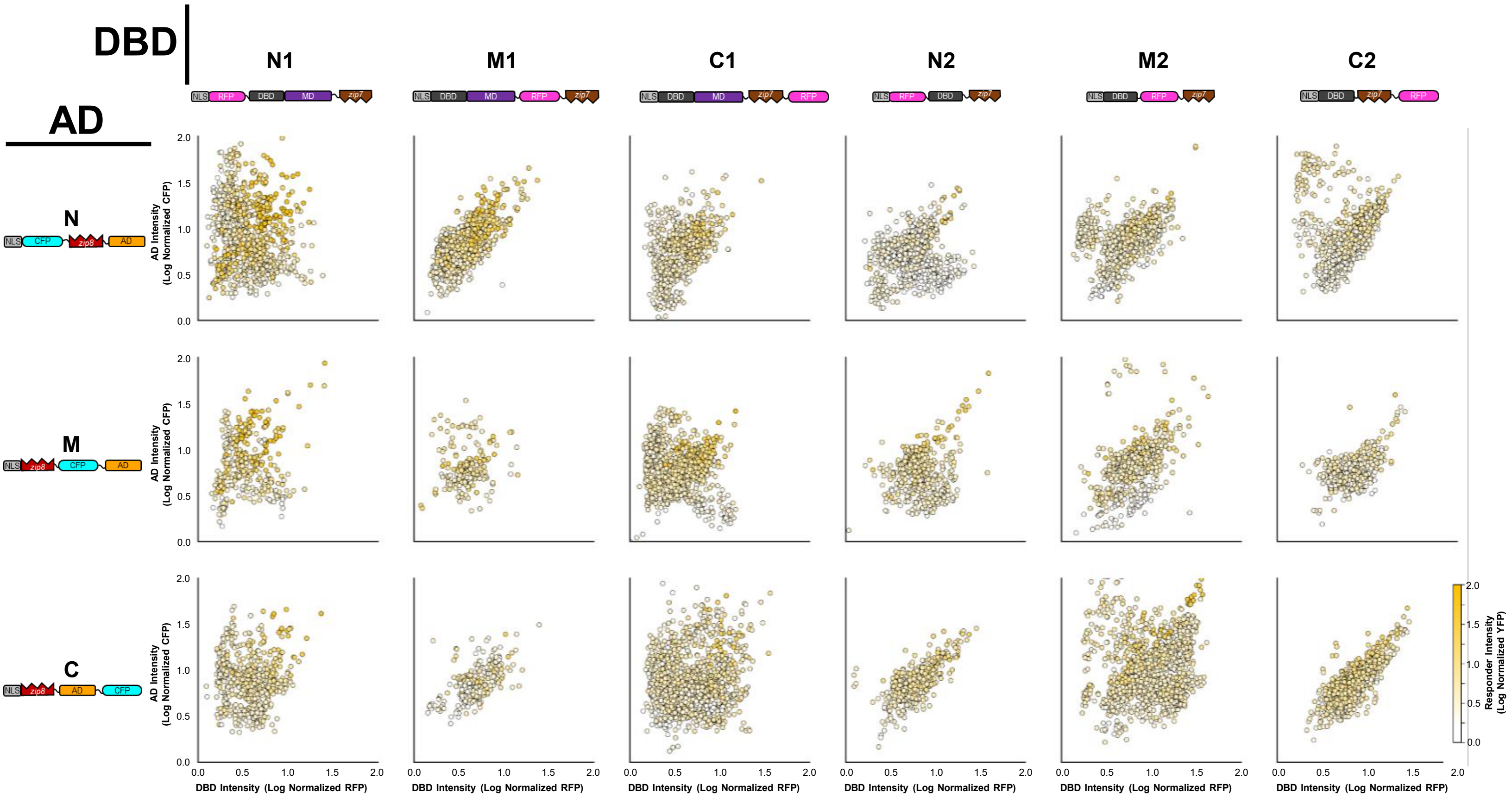

### Supplemental Figure 4: Tagged hemidriver topology experiments all decile plots

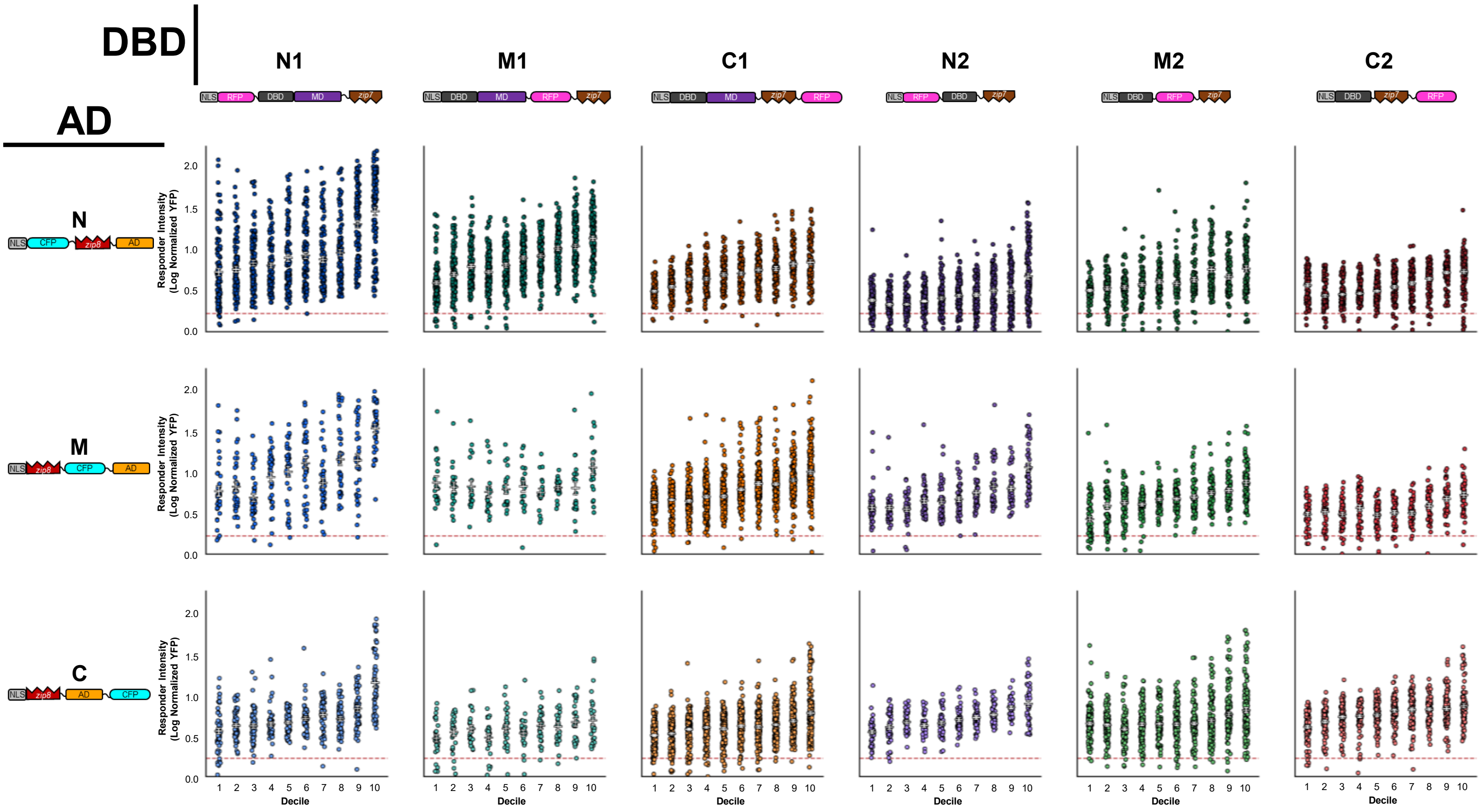

### Supplemental Figure 5: Replacement of fluorescent protein tag on hemidriver with HaloTag

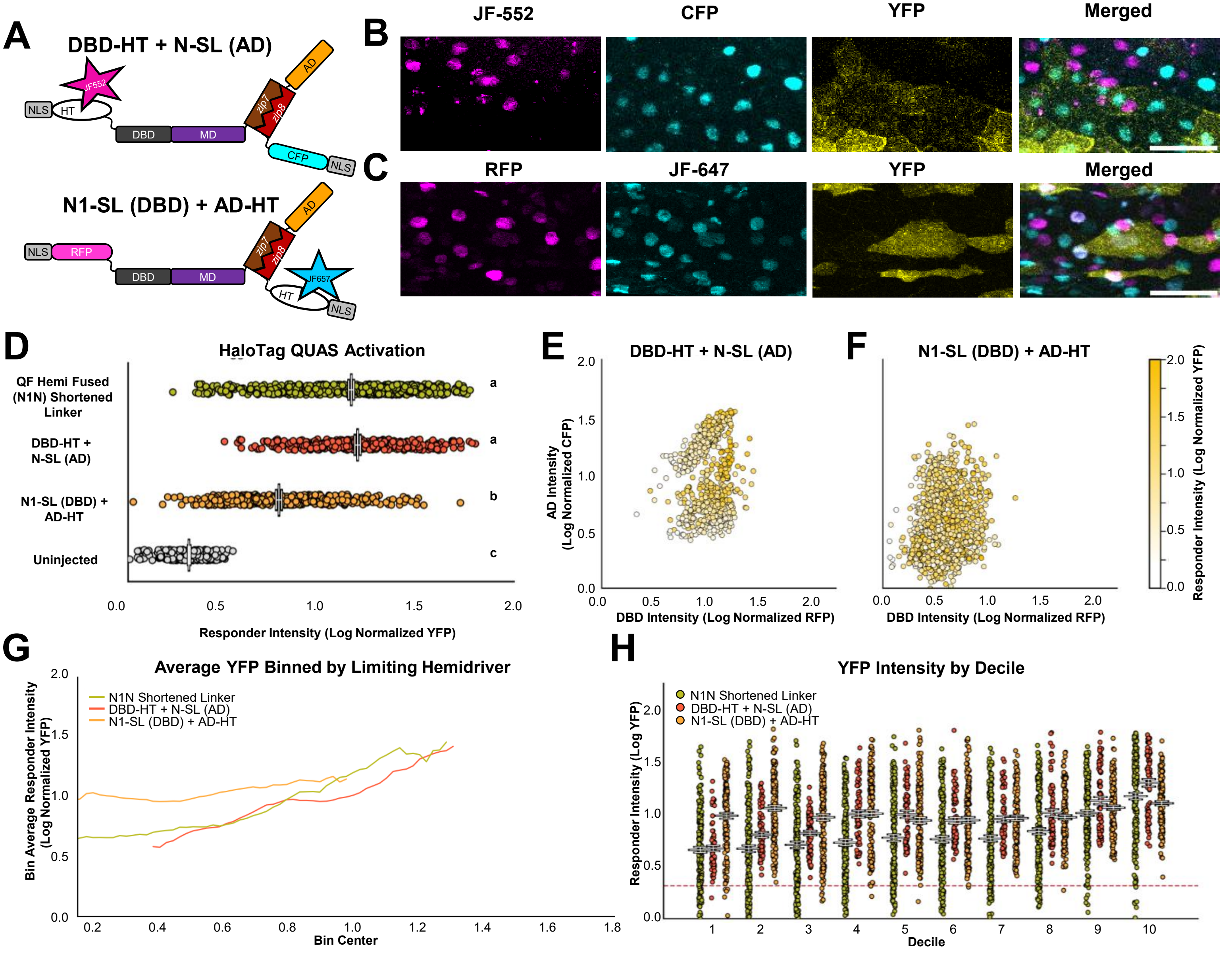

### Supplemental Figure 6: Individual P2A hemidriver solo injections

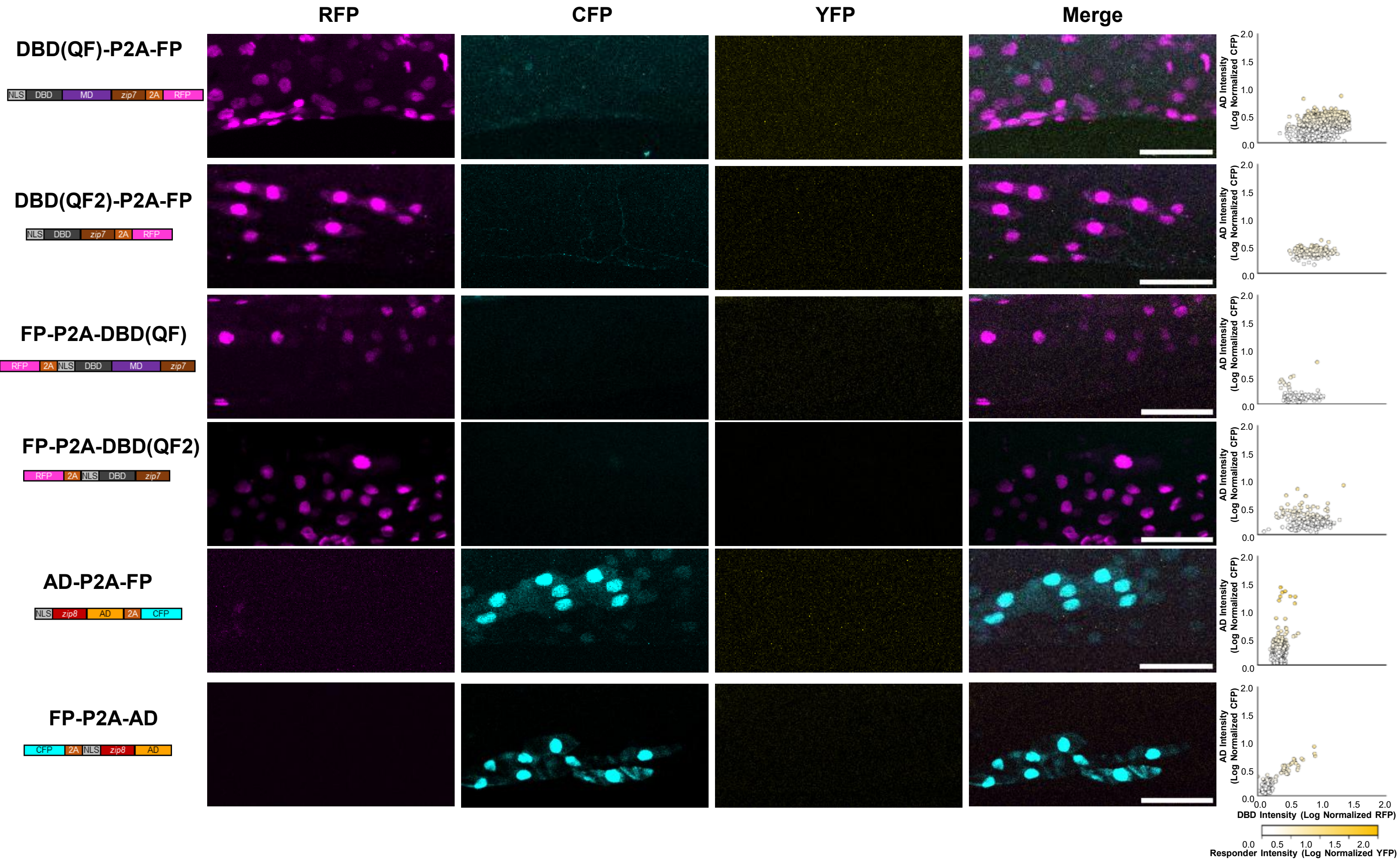
