## Supplemental File 14 for "Fluorescently-labeled split-QF hemidrivers: simplifying and enhancing methods enabling intersectional targeting of discrete cell types"

Supplemental File: Suggested Hemidrivers for Specific Applications

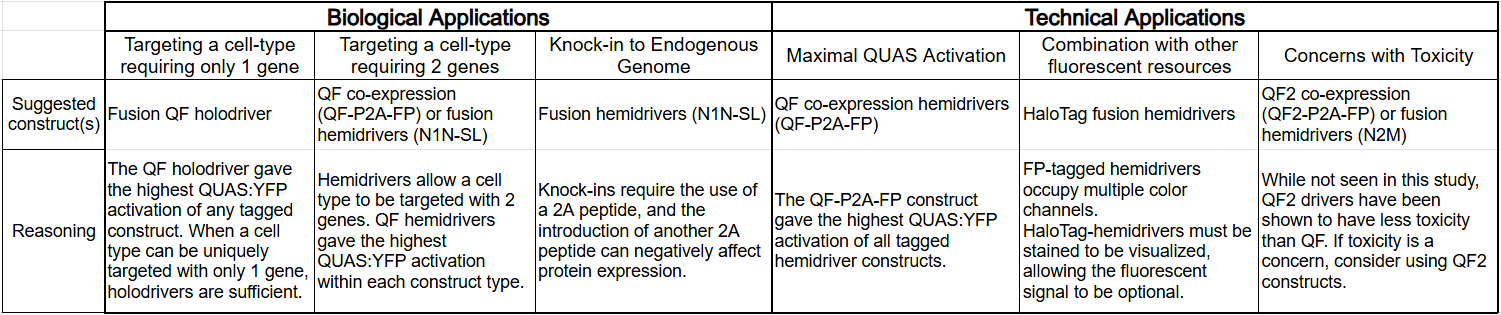

Table Legend:

Additional factors must be considered when selecting among labeled hemidriver options for a specific purpose. For instance, using 2A elements for co-expressing reporters resulted in the highest degree of hemidriver transactivation. However, if multiple 2A sequences are required (e.g., for non-disruptive knock in constructs) the possibility of unpredictable effects on gene expression arises^1^. In this case, additional controls would be needed to ensure expression fidelity is retained for the targeted gene and intersectional components^1^. Other strategies for polycistronic expression (such as the internal ribosome entry site, IRES) also face significant technical limitations, including inefficient expression of downstream elements, relatively long and complex nucleotide sequences, and declining activity when arrayed in tandem. Therefore, despite having a lower QUAS activation level, fusion hemidrivers would be better suited for knocking into an endogenous gene. Another situation that warrants caution is the use of hemidriver FP fusions/co-expression in the presence of other transgenes with fluorophores. Tagging hemidrivers affords many advantages, but it uses multiple color channels and limits the ability to combine them with other fluorescent transgenes or antibodies. In these situations, we recommend the use of self-labeling protein tags, such as HaloTag, to provide the option for the hemidrivers to be fluorescently labeled or unlabeled. Thus, the hemidriver-HaloTag construct can be irreversibly stained and visualized in select samples, retaining the advantages of tagging, while other samples can be left unstained, leaving those color channels available for additional transgenes. These self-labeling tags permit the possibility of combining resources with other self-labeling proteins, such as CLIP-tag or SNAP-tag. Of note, we could not successfully stain and visualize the SNAP-tag construct, despite observing QUAS activation (data not shown), perhaps because it stains much slower and dimmer than HaloTag^2^. Fortunately, a newer version, SNAP-tag2, was recently released and produces faster and brighter labeling than the original SNAP-tag^3^, giving promise for use in combination with other self-labeling protein constructs, such as the HaloTag hemidriver constructs. Accordingly, we recommend selecting hemidriver variants depending on the precise application.
